# Hypoxia delays steroid-induced developmental maturation in *Drosophila* by suppressing EGF signaling

**DOI:** 10.1101/2023.05.12.540517

**Authors:** Michael J Turingan, Tan Li, Jenna Wright, Savraj S Grewal

## Abstract

Animals often grow and develop in unpredictable environments where factors like food availability, temperature, and oxygen levels can fluctuate dramatically. To ensure proper sexual maturation into adulthood, juvenile animals need to adapt their growth and developmental rates to these fluctuating environmental conditions. Failure to do so can result in impaired maturation and incorrect body size. Here we describe a mechanism by which *Drosophila* larvae adapt their development in low oxygen (hypoxia). During normal development, larvae grow and increase in mass until they reach critical weight (CW), after which point a neuroendocrine circuit triggers the production of the steroid hormone ecdysone from the prothoracic gland (PG), which promotes maturation to the pupal stage. However, when raised in hypoxia (5% oxygen), larvae slow their growth and delay their maturation to the pupal stage. We find that, although hypoxia delays the attainment of CW, the maturation delay occurs mainly because of hypoxia acting late in development to suppress ecdysone production. This suppression operates through a distinct mechanism from nutrient deprivation, occurs independently of HIF-1 alpha and does not involve modulation of PTTH, the main neuropeptide that initiates ecdysone production in the PG. Instead, we find that hypoxia lowers the expression of the EGF ligand, spitz, and that the delay in maturation occurs due to reduced EGFR/ERK signaling in the PG. Our study sheds light on how animals can adjust their development rate in response to changing oxygen levels in their environment. Given that hypoxia is a feature of both normal physiology and many diseases, our findings have important implications for understanding how low oxygen levels may impact animal development in both normal and pathological situations.

## Introduction

Animals often grow and develop in unpredictable environments. Fluctuations in nutrient availability, temperature, or oxygen and exposure to toxins and pathogens can create conditions of environmental stress. Animals must, therefore, be able to assess these conditions and, in turn, adapt their physiology and growth rate to ensure proper development and attainment of functional organ and body size (Callier et al., 2015; Callier and Nijhout, 2014; Koyama and Mirth, 2018; Koyama et al., 2020; Mirth et al., 2021; Troha and Ayres, 2020; Wang et al., 2006). Any defects in these adaptations can impair fitness. Understanding how environmental signals control animal development is, therefore, an important question in biology.

In animals that exhibit determinate growth, increases in body size occur during the juvenile stage of development before they mature to an adult stage, and further growth stops. Final body size is thus determined by the duration of the juvenile growth stage and the timing of maturation. *Drosophila* has provided a tractable and valuable model system to understand how environmental conditions can control the timing of juvenile maturation (Rewitz et al., 2013; Yamanaka et al., 2013). In *Drosophila*, growth occurs during the larval (juvenile) period where animals increase ∼200-fold in mass over a 4-5 day period before undergoing maturation to the pupal stage and metamorphosis into adults (Britton and Edgar, 1998; Church and Robertson, 1966). Since adults do not grow, the timing of larval maturation is a key determinant of final body size. This timing is controlled by a pulse of secretion of the steroid hormone ecdysone from the prothoracic gland (PG), which then acts on larval tissues to promote the transition to the pupal stage (Yamanaka *et al*., 2013). Two subsets of neurons control this ecdysone pulse – a bilateral pair of PTTH neurons and a bilateral subset of serotonergic neurons – that both innervate the PG and stimulate ecdysone production (McBrayer et al., 2007; Shimada-Niwa and Niwa, 2014; Shimell et al., 2018). In addition, several PG-derived autocrine factors and endocrine signals from other tissues can act on the PG to control ecdysone synthesis through multiple cell-cell signalling pathways (Kannangara et al., 2021; Malita and Rewitz, 2021; Pan et al., 2021). Together, these mechanisms provide a way for the PG to integrate various cues to ensure the proper timing of ecdysone production and triggering of larval maturation (Danielsen et al., 2013; Di Cara and King-Jones, 2016).

One cue that is perhaps the best-understood environmental regulator of larval maturation is nutrient availability (Danielsen *et al*., 2013; Malita and Rewitz, 2021). For larval maturation to occur properly, larvae need to attain an appropriate mass and level of nutrient stores - defined as critical weight (CW) - to support metamorphosis during the pupal stage (Malita and Rewitz, 2021). Hence, larvae must be able to sense their nutritional status and then adapt their development rate accordingly – in rich nutrient condition larval development proceeds rapidly, but when nutrients are scarce larval development and maturation are delayed. The conserved insulin and TOR kinase signaling pathways mediate nutrient sensing and signaling in larvae (Boulan et al., 2015; Grewal, 2009; Hietakangas and Cohen, 2009; Texada et al., 2020), and several studies have shown that stimulation of insulin/TOR signaling in the PG provides a mechanism to couple nutrients to CW (Colombani et al., 2005; Ghosh et al., 2022; Koyama et al., 2014; Layalle et al., 2008; Mirth et al., 2005; Ohhara et al., 2017; Zeng et al., 2020). In addition, the serotonergic neurons that innervate the PG can link nutrient availability to the production of ecdysone in the PG (Shimada-Niwa and Niwa, 2014), while PTTH has been shown to be important for allowing larvae to adapt the timing of their maturation to poor nutrient and crowded growth conditions (Shimell *et al*., 2018).

Oxygen availability is another key environmental cue that can modulate animal development (Bickler and Buck, 2007; Ducsay et al., 2018; Moore et al., 2011; Zhou and Haddad, 2013). *Drosophila* larvae grow and develop by burrowing into fermenting food rich in microorganisms, an environment that is likely characterized by low oxygen or hypoxia (Callier *et al*., 2015; Markow, 2015). Hence, they have evolved mechanisms to be able to adapt their development to conditions of hypoxia. When grown in moderate hypoxia (5-10% oxygen) larvae slow their growth rate and delay their maturation leading to smaller sized adults (Callier *et al*., 2015; Callier and Nijhout, 2014; Callier et al., 2013; Farzin et al., 2014; Harrison et al., 2018; Harrison and Haddad, 2011; Kapali et al., 2022; Lee et al., 2019; Peck and Maddrell, 2005; Texada et al., 2019; Wong et al., 2014). Studies have described how the reduction in growth rate occurs through reduced insulin and TOR kinase signaling (Barretto et al., 2020; Kapali *et al*., 2022; Lee *et al*., 2019; Texada *et al*., 2019; Wong *et al*., 2014). However, the mechanisms by which hypoxia delays the timing of larval maturation remain to be determined. We explore this issue in this study.

## Results

### Hypoxia slows development and delays larval maturation

We raised larvae in normoxia or hypoxia (5% oxygen) from hatching and measured the time to pupation. We saw that the hypoxia exposed larvae showed an approximate two-day delay in development (Figure 1A). As previously described (Lee *et al*., 2019; Texada *et al*., 2019), this was accompanied by a reduction in final body size but without any reduction in larval viability (Figure S1A, B). These results show that hypoxia exposed larvae have reduced body growth and a delay in development to the pupal stage. Larval maturation is regulated by a neuroendocrine circuit that produces a pulse of ecdysone from the prothoracic gland (PG) which triggers the larval to pupal transition (Yamanaka *et al*., 2013). To measure this ecdysone pulse, we used qPCR to quantify mRNA levels of two Halloween genes, *phm* and *spok*, that are involved in ecdysone biosynthesis in the PG. We found that, in normoxia, the expression of these genes peaked at 144 hr AEL (after egg laying), prior to the larval to pupal transition. However, in the hypoxia-exposed larvae the increase in expression of both *phm* and *spok* were delayed and blunted compared to their normoxic counterparts (Figure 1B). These results indicate that the hypoxia-induced delay in larval maturation is associated with both a delay and reduction in the final larval ecdysone pulse.

**Figure 1.**
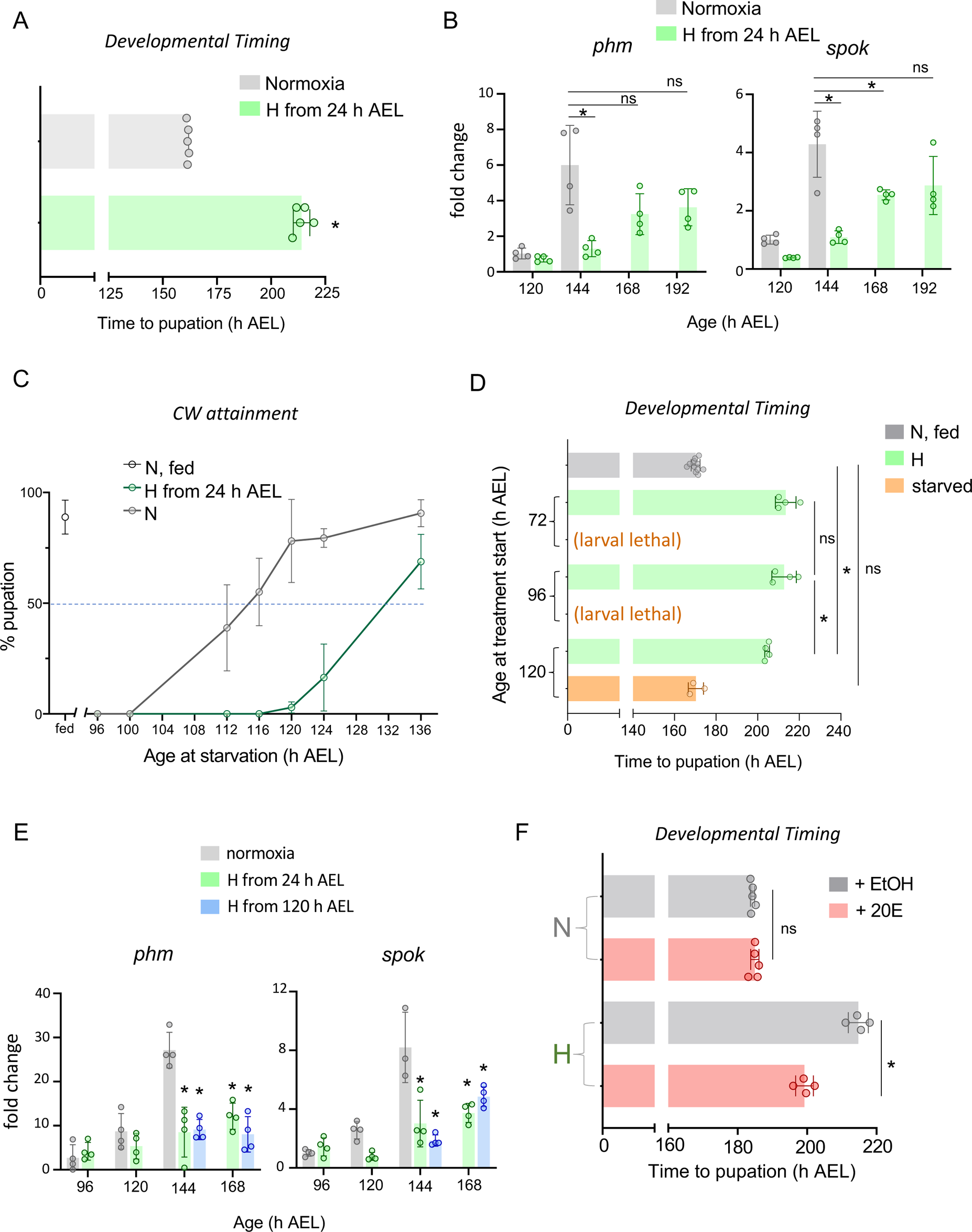
Hypoxia can delay ecdysone production and larval maturation independent of critical weight attainment. **(A)** Average time to pupation of *w^1118^* larvae reared in ambient oxygen or 5% O_2_ (from hatching), in hours after egg laying (h AEL). Each data point represents a vial of 30 larvae. n (# of vials of 30) ≥ 3 per condition. **(B)** Relative mRNA levels (normalized to Rpl32) of ecdysone biosynthetic genes *phantom* (*phm*) and *spookier* (*spok*) from whole-larvae qRT-PCR of *w^1118^* larvae reared in ambient oxygen or in 5% O2 from hatching (24 h AEL). n (# of independent samples) ≥ 3 per condition. **(C)** Determination of age at Critical weight (CW) attainment for animals reared in ambient or 5% O_2_ (Hypoxia, ‘H’) conditions. The age at CW attainment (in h AEL) was defined as the age which yielded 50% pupation upon starvation. Each data point represents the average of three vials of 30 larvae. n (# of vials of 30) ≥ 3 per condition. **(D)** Developmental timing of *w^1118^ larvae* either starved or exposed to 5% O_2_ at the indicated larval age. n (# of vials of 30) ≥ 3 per condition **(E)** Relative mRNA levels (normalized to Rpl32) of ecdysone biosynthetic genes *phantom* (*phm*) and *spookier* (*spok*) from whole-larvae qRT-PCR of *w^1118^* larvae reared in ambient oxygen or in 5% O_2_ from either 24 h AEL or 120 h AEL. n (# of independent samples) ≥ 3 per condition. **(F)** Developmental timing of *w^1118^* larvae fed either 20-hydroxyecdysone (20E) or ethanol (EtOH) vehicle, reared in either ambient oxygen or 5% oxygen from 120 h AEL (*i.e.,* post-CW). n (# of vials of 30) ≥ 3 per condition. * denotes p < 0.05. Also see Fig S1.

### Hypoxia acts post critical weight to suppress ecdysone signaling

A key checkpoint for the initiation of larval maturation is the attainment of critical weight (CW), a point in larval development where animals have achieved sufficient mass and nutrient stores to complete development to viable adults without any further nutrients (Danielsen *et al*., 2013; Rewitz *et al*., 2013). CW is determined, in large part, by an animal’s growth rate, which in *Drosophila* larvae is controlled by the conserved insulin and TOR kinase signaling pathways (Malita and Rewitz, 2021). Given that hypoxia exposure can suppress larval growth by inhibiting these pathways (Barretto *et al*., 2020; Kapali *et al*., 2022; Lee *et al*., 2019; Texada *et al*., 2019; Wong *et al*., 2014), it is possible that the delay in maturation merely reflects a hypoxia-induced delay in attainment of CW. To address this, we raised larvae from hatching in either normoxia or hypoxia and then at defined times in development we switched them to starvation conditions and monitored their subsequent development in normoxia. CW can be assessed by determining the time point in development at which >50% larvae can pupariate when subsequently starved. We found that in normoxia this point was reached at ∼114 hrs AEL, whereas in hypoxia-exposed animals, this point was at ∼132hrs AEL, indicating that hypoxia exposure does delay CW (Figure 1C). We then examined the effects of hypoxia exposure introduced either before or after CW attainment. We raised larvae from hatching in normoxia and then at 72hrs, 96 hrs or 120 hrs AEL we switched larvae either to hypoxia or starvation conditions and then monitored their subsequent development. As expected, we found that larvae starved at 72 and 96hrs (pre-CW) arrested their development and failed to pupate while larvae starved at 120 hrs (post CW) completed larval development and pupated at the same time as control (normoxic, fed) larvae but at a significantly smaller final size (Figure 1D and Figure S1C, D). However, we found that larvae exposed to hypoxia at all three time points were able to pupate but showed a delay to pupation (Figure 1D and Figure S1C). And, interestingly, larvae switched to hypoxia post CW (120hrs AEL) showed a delay in development (∼34hrs) that was only slightly shorter than that seen with pre-CW larvae switched to hypoxia at 72hrs AEL (43 hr delay) or 96hrs AEL (42 hrs delay) (Figure 1D). We also found that animals switched to hypoxia exposure at each of three timepoints had significantly reduced pupal sizes compared to normoxic controls, but that animals exposed to hypoxia post-CW were significantly larger than those exposed to hypoxia pre-CW (Figure S1D). Together these results indicate that, although hypoxia early in larval development can slow growth and delay CW attainment, hypoxia can also act specifically at the late larval stage to suppress the mechanism(s) that promote maturation. Consistent with this we saw that larvae exposed to hypoxia post CW (120hrs AEL) showed a similar suppression of *phm* and *spok* mRNA levels as larvae exposed to hypoxia from hatching (Figure 1E), suggesting that the hypoxia-induced decrease in ecdysone biosynthesis is largely due to an effect of low oxygen in late larval development. Furthermore, we found that feeding larvae 20-hydroxyecdysone could reverse the delay in development seen in larvae that were switched to hypoxia post CW (at 120hrs AEL) and led to smaller sized animals compared to vehicle-fed hypoxic animals (Figure 1F and Figure S1E). These results suggest that hypoxia can act on the mechanisms that control the timing of the late larval ecdysone pulse. They also suggest that larvae may use different mechanisms to couple nutrient availability and oxygen levels to the control of maturation timing. Previous work has described how stimulation of TOR kinase activity in the PG is a key mechanism that couple nutrient availability to the attainment of CW and larval maturation, and that genetic activation of TOR signaling in the PG can reverse the delay in larval development see in low nutrient conditions(Layalle *et al*., 2008). However, we found that overexpression of the TOR activator, Rheb, in the PG could not reverse the hypoxia-mediated delay in development (Figure S1F), providing further evidence that nutrient restriction and hypoxia may operate through different mechanisms to delay larval maturation.

### The hypoxia maturation delay is independent of HIF1-alpha/sima function in neurons or the PG

The conserved hypoxia-inducible factor 1 alpha (HIF-1 alpha), known as *sima* in *Drosophila*, is perhaps the best characterized mediator of hypoxia responses in metazoans (Semenza, 2011). In normoxia, HIF-1 alpha protein is continually synthesized but then rapidly degraded. However, upon hypoxia exposure this degradation is blocked, HIF-1 alpha protein accumulates, and it can then translocate to the nucleus and control the expression of genes needed for hypoxic adaptive responses (Semenza, 2011). Previous studies have shown that *sima* can modulate growth and survival during periods of hypoxia in flies (Centanin et al., 2005; Lavista-Llanos et al., 2002; Li et al., 2013; Romero et al., 2007; Texada *et al*., 2019). We therefore explored whether *sima* might mediate the maturation delay seen in hypoxia. Given the central role for the PG in controlling ecdysone production and larval maturation, we first examined the effects of RNAi-mediated knockdown of *sima* in the PG. We found that RNAi-knockdown of *sima* using the PG driver, *P0206-Gal4,* led to a modest delay in development in normoxia but did not reverse the hypoxia-induced delay in the larval-to-pupal transition (Figure 2A). Knockdown of *sima* with another PG driver, *phm-Gal4*, also did not lead to a reversal in the hypoxia-induced delay in larval maturation (Figure 2B). Given the central role in neuronal signaling to the PG in controlling ecdysone production, we also examined a role for neuronal *sima* in the hypoxia-mediated delay in maturation. We used the *elav-Gal4* line to drive *UAS-sima RNAi* specifically in all post-mitotic neurons. We found that neuronal-specific knockdown of *sima* had no effect on the timing of larval maturation in animals grown in either normoxia or hypoxia (Figure 2C). Similar results were seen when we used a second independent *UAS-Sima RNAi* line (Figure S2). Together, these results suggest that the hypoxia-mediated delay in larval maturation does not rely on induction of HIF1-alpha/sima in the neuron-PG neuroendocrine network responsible for stimulating ecdysone production.

**Figure 2.**
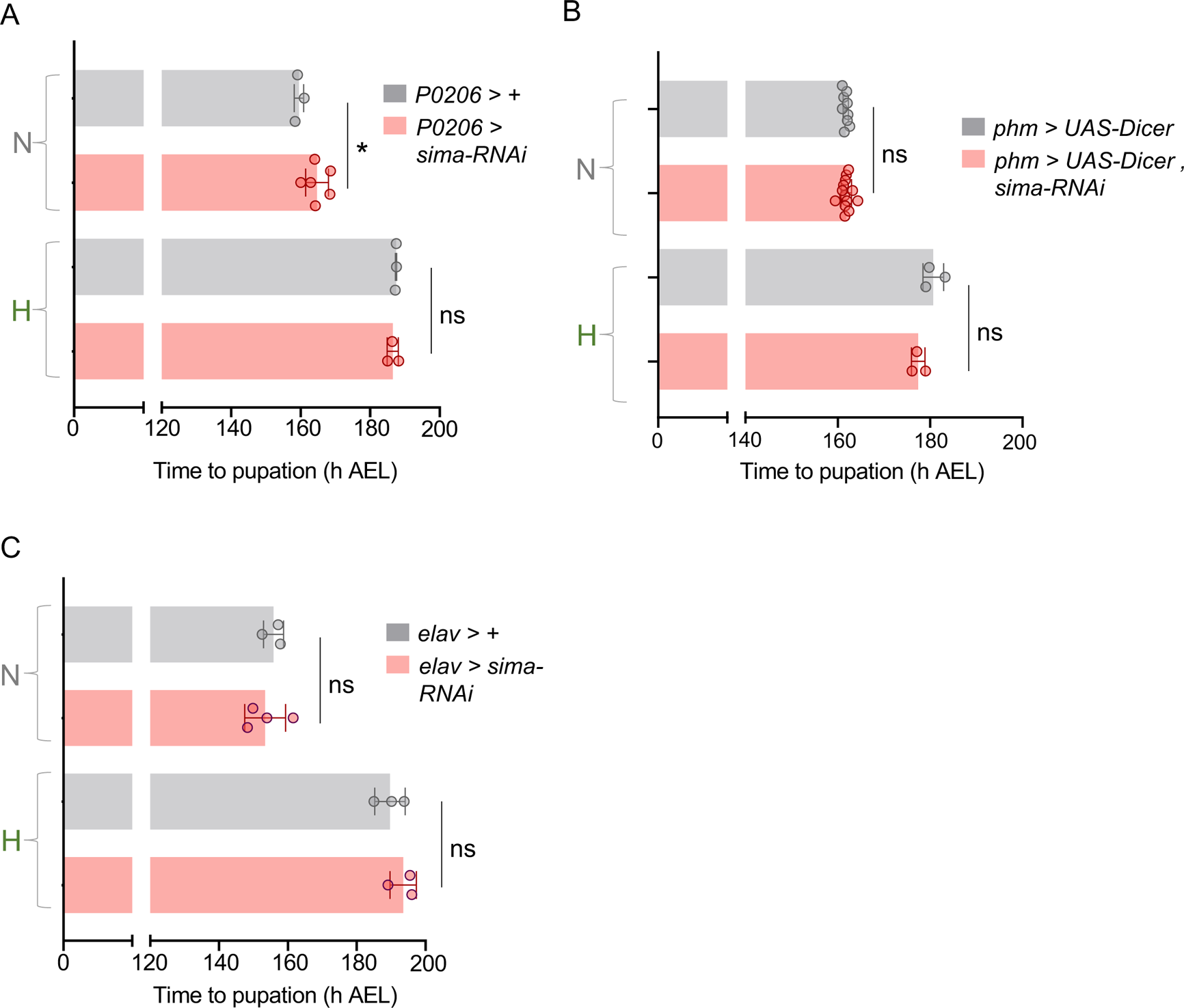
The hypoxia-induced developmental delay is independent of sima/HIF-1α. **(A-C)** Average time to pupation of larvae of the indicated genotype reared in either normal oxygen conditions throughout development (‘N’) or shifted to 5% O_2_ at 120 h AEL (‘H’). Knockdown of *sima* was performed in the prothoracic gland/corpus allatum (*P0206-GAL4*) **(A)**, prothoracic gland alone (*phm-GAL4, UAS-dicer*) **(B)**, or pan-neuronally (*elav-GAL4*) **(C)**. n (# of vials of 30) ≥ 3 per condition. * denotes p < 0.05. Also see Fig S2.

### The hypoxia maturation delay is independent of PTTH/Torso signaling

A key regulator of ecdysone production is the neuropeptide PTTH, which is secreted from a pair of PG-innervating neurons (the ptth neurons) and then binds to the receptor tyrosine kinase (RTK) Torso on PG cells to trigger ecdysone synthesis (Rewitz et al., 2009; Yamanaka *et al*., 2013). We therefore explored the effects of manipulating PTTH signaling on the hypoxia-mediated delay in larval maturation. We first examined *ptth* mutants. As described previously we found that, in normoxia, *ptth* null mutants showed a delay in pupation and a 13% increase in final pupal size (Figure 3A, B) (McBrayer *et al*., 2007; Shimell *et al*., 2018), consistent with a role for the PTTH signaling in the control of timing the late larval ecdysone pulse. However, we found that this delay in maturation was further exacerbated in hypoxia-exposed larvae (Figure 3A) and *ptth* mutants now showed a 24% increase in final pupal size (Figure 3B). We saw similar effects with another *ptth* mutant (Figure S3A, B). We also examined the effects of knockdown of the PTTH receptor, Torso, in the PG. We found that RNAi-mediated knockdown of *torso* in the PG of normoxic animals led to a delay in pupation and a 24% increase in pupal size. However, as with loss of PTTH, the delay in development seen with *torso* knockdown was further exacerbated in hypoxia exposed animals (Figure 3C) and *torso* knockdown animals now showed a 31% increase in pupal size. Together, these additive effects of hypoxia and loss of PTTH/Torso signaling on larval maturation suggest that the effects of hypoxia on larval maturation occur independently of suppression of PTTH/Torso signaling.

**Figure 3.**
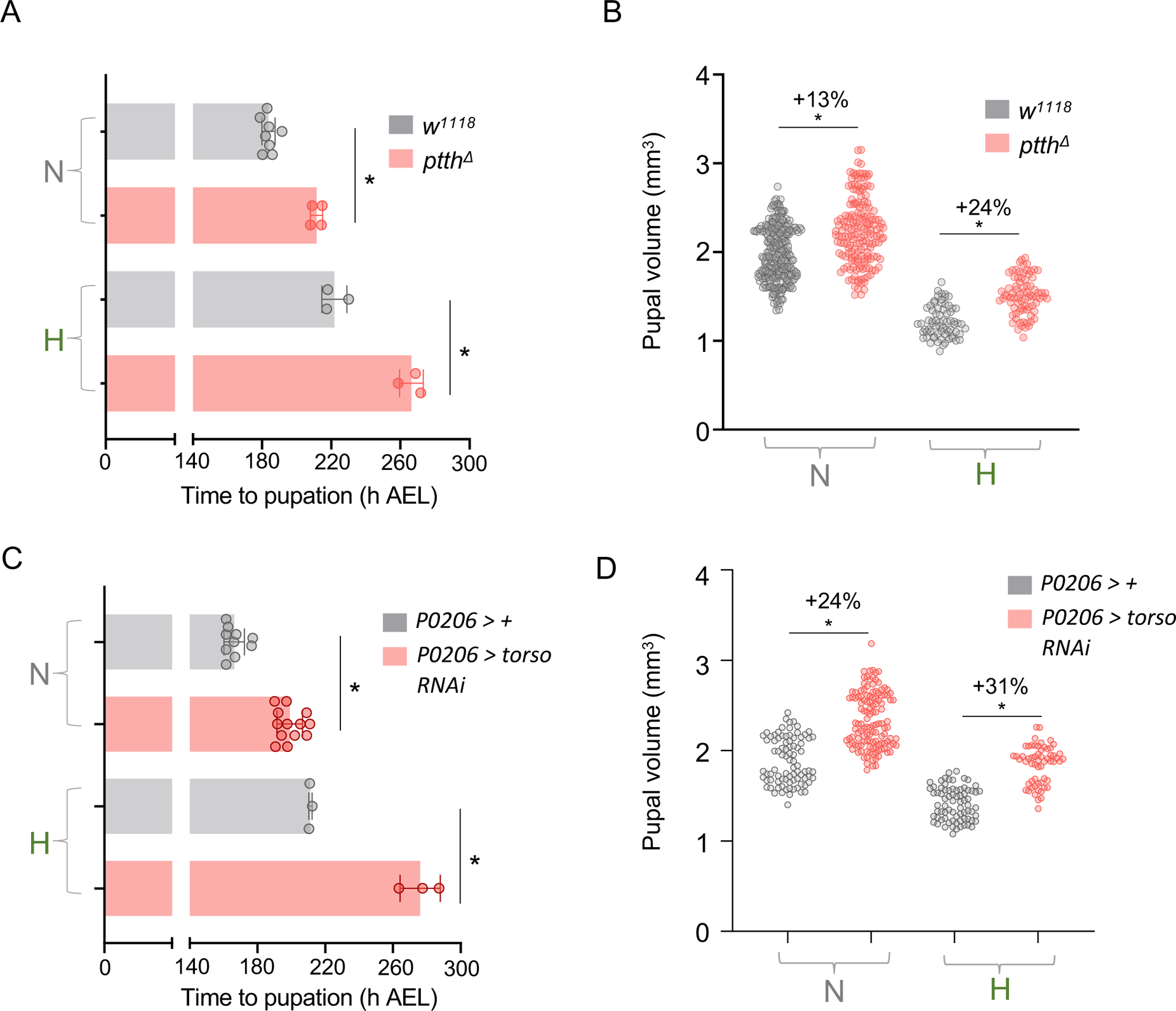
The hypoxia-induced maturation delay is independent of PTTH/Torso signaling. **(A)** Average time to pupation of larvae, either *w^1118^* or mutant for *ptth*, reared in either normal oxygen conditions throughout development (‘N’) or shifted to 5% O_2_ at 120 h AEL (‘H’) n (# of vials of 30 larvae) ≥ 3 per condition. **(B)** Pupal size of animals, either *w^1118^* or mutant for *ptth*, reared in normoxia or hypoxia from 120 h AEL. Each data point represents body size measured for one animal. n (# of pupae) = 254 (N *w^1118^*), 172 (N *ptth^Δ^*), 70 (H *w^1118^*), 79 (H *ptth^Δ^*). **(C)** Average time to pupation of larvae, either *P0206 >* + or *P0206 > torso RNAi*, reared in either normal oxygen conditions throughout development or shifted to 5% O_2_ at 120 h AEL. n (# of vials of 30 larvae) ≥ 3 per condition. (D) Pupal size of animals, either *P0206 >* + or *P0206 > torso RNAi,* reared in normoxia or hypoxia from 120 h AEL. Each data point represents body size measured for one animal. n (# of pupae) = 86 (N *P0206 >* +), 127 (N *P0206 > torso RNAi*), 75 (H *P0206 >* +), 59 (H *P0206 > torso RNAi*). * denotes p < 0.05. Also see Fig S3.

### Hypoxia induces a delay in maturation by suppressing spitz-EGFR signaling in the PG

Stimulation of the Ras/ERK signaling pathway in the PG is the key trigger for inducing the late larval pulse of ecdysone synthesis required for pupation (Yamanaka *et al*., 2013). We therefore examined the effects of manipulating Ras/ERK signaling in the PG on the hypoxia-induced delay in larval maturation. We found that PG expression (using *P0206-Gal4*) of an activated form of Raf kinase (*UAS-Raf^GOF^*) led to pronounced acceleration in the time to pupation in normoxic larvae (Figure 4A). This result is consistent with previous work showing that genetic activation of Ras/ERK signaling in the PG can induce precocious pupation by accelerating the onset of the late larval ecdysone pulse (Rewitz *et al*., 2009). Interestingly, we found that PG expression of *UAS-Raf^GOF^* using *P0206-Gal4* also accelerated the time to pupation in hypoxic larvae (Figure 4A). This accelerated development was even more pronounced when we expressed *Raf^GOF^* with a different PG driver (*spok-Gal4)* and, in this case, the hypoxia-mediated delay in development was almost completely abolished with hypoxia exposed animals pupating only slightly later than normoxic animals (Figure S4A).

**Figure 4.**
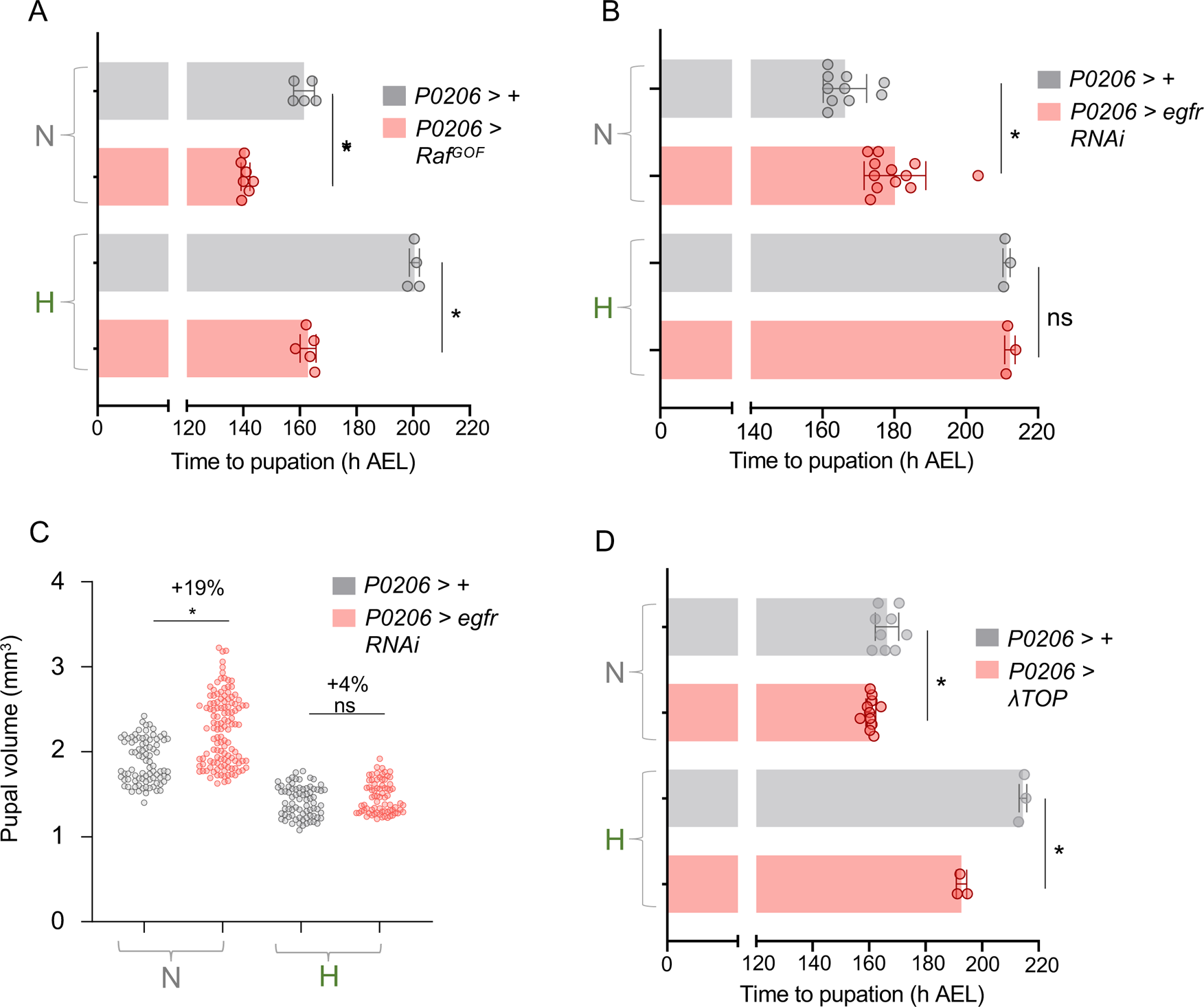
Hypoxia induces a maturation delay by suppressing Egfr/MAPK signaling. **(A)** Average time to pupation of larvae, either *P0206 >* + or *P0206 > Raf^GOF^*, reared in either normal oxygen conditions throughout development (‘N’) or shifted to 5% O_2_ at 120 h AEL (‘H’). n (# of vials of 30 larvae) ≥ 3 per condition. **(B)** Average time to pupation of larvae, either *P0206 >* + or *P0206 > egfr-RNAi*, reared in either normal oxygen conditions throughout development or shifted to 5% O_2_ at 120 h AEL. n (# of vials of 30 larvae) ≥ 3 per condition. **(C)** Pupal size of animals, either *P0206 >* + or *P0206 > egfr-RNAi,* reared in normoxia or hypoxia from 120 h AEL. Each data point represents body size measured for one animal. n (# of pupae) = 86 (N *P0206 >* +), 123 (N *P0206 > egfr-RNAi*), 75 (H *P0206 >* +), 74 (H *P0206 > egfr-RNAi*). **(D)** Average time to pupation of larvae, either *P0206 >* + or *P0206 > λTOP*, reared in either normal oxygen conditions throughout development or shifted to 5% O_2_ at 120 h AEL. n (# of vials of 30 larvae) ≥ 3 per condition. * denotes p < 0.05. Also see Fig S4.

A recent paper showed that signalling through Epidermal Growth Factor RTK (EGFR) is the main contributor to Ras/ERK stimulation in the PG (Cruz et al., 2020). Hence, we explored whether modulation of EGFR affects the hypoxia-mediated delay in pupation. We found that RNAi-mediated knockdown of EGFR using *phm-Gal4* prevented pupation, as previously reported (Figure S4B), making it difficult to assess effects of hypoxia on pupation in these larvae. However, we saw that EGFR knockdown with an alternate PG driver, *P0206-Gal4*, led to a delay in development in normoxic animals and an increase in final pupal size, consistent with the role for EGFR signaling in the timing of maturation (Figure 4B, C). Interestingly, this delay to pupation was not further exacerbated by hypoxia exposure (Figure 4B) and the EGFR knockdown animals raised in hypoxia showed no significant difference in final pupal size compared to controls (Figure 4C). This lack of additive effects of EGFR knockdown and hypoxia on delayed development suggest that hypoxia may delay larval maturation by suppressing EGFR signaling. To test this further we examined the effects of overexpression of a constitutively active form of the EGFR in the PG and found that this led to a partial reversal of the hypoxia-mediated delay in development (Figure 4D). Together, these results suggest that one way that hypoxia delays larval maturation is by decreasing EGFR-mediated signalling in the PG. Signalling through two other RTKs, Alk and Pvr, in the PG has also been shown to contribute to Ras/ERK activation and the control of ecdysone synthesis and developmental timing (Juarez-Carreño and Geissmann, 2023; Pan and O’Connor, 2021). However, we found that RNAi-mediated knockdown of either of these RTKs had little effect on developmental timing (Figure S4C, D), suggesting that suppression of either pathway is unlikely to contribute to the hypoxia-induced delay in larval maturation.

There are two main ligands, vein and spitz, that stimulate EGFR in the PG to promote maturation (Cruz *et al*., 2020). Interestingly, we saw an upregulation of *spitz* mRNA in late third instar normoxic larvae, as they approach pupation, and this increased expression was suppressed in larvae that were either raised in hypoxia from hatching or switched to hypoxia post-CW (at 120 hrs)(Figure 5A). In normal development, PG-derived Spitz functions in an autocrine manner to stimulate the EGFR-dependent control of larval maturation. We therefore specifically examined whether hypoxia suppresses Spitz levels in the PG. To do this, we made use of a *spitz-LacZ* enhancer trap line that expresses beta-galactosidase under the control of the *spitz* gene promoter. Using this line, we found that PG LacZ levels were reduced in larvae switched to hypoxia post-CW (at 120 hrs) compared to normoxic age-matched controls (Figure 5B). These results suggest that one way that hypoxia might suppress ecdysone production in the PG is by preventing the late larval stage upregulation of spitz-induced EGFR signaling. We therefore examined the effects of manipulating PG Spitz levels on the hypoxia-induced delay in maturation. We found that RNAi-mediated knockdown of Spitz in the PG delayed pupation in normoxic animals, as described before (Figure 5C). However, PG-specific spitz knockdown did not add to the hypoxia-mediated delay in larval maturation, suggesting that this delay may rely on reduced Spitz signalling (Figure 5C). To test this further, we examined the effects of overexpression of Spitz in the PG. We found that PG-specific overexpression of Spitz had little effect on developmental timing in normoxia (Figure 5D). In contrast, Spitz overexpression led to a strong reversal of the hypoxia-induced developmental delay to a similar extent as that seen with 20E feeding (Figure 5D).

**Figure 5.**
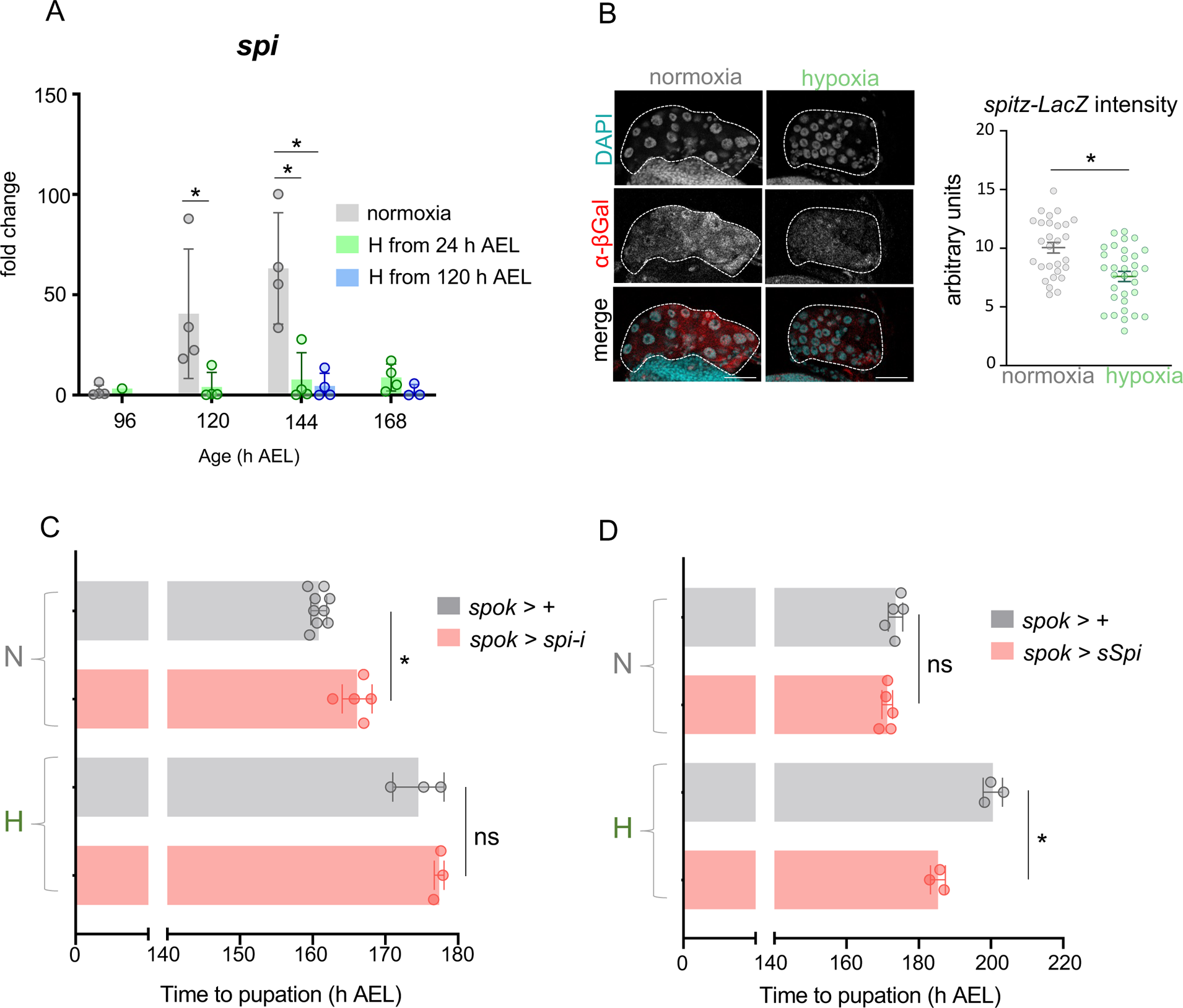
Hypoxia supresses spitz expression to delay Egfr/MAPK-dependent maturation. **(A)** Relative mRNA levels (normalized to Rpl32) of Egf ligand *spitz* from whole-larvae qRT-PCR of *w^1118^* larvae reared in ambient oxygen or in 5% O_2_ from either 24 h AEL or 120 h AEL. n (# of independent samples) ≥ 3 per condition, except for hypoxic larvae at 96 h AEL. **(B)** Representative confocal micrographs and immunofluorescence quantification for anti-beta-galactosidase immunostaining of PGs (indicated with dashed outline) from 140 h AEL *spitz-LacZ* larvae, either reared in normoxia throughout development or in hypoxia from 120 h AEL. Scale bar indicates 50 μm. **(C)** Average time to pupation of larvae, either *spok >* + or *spok > spitz-RNAi*, reared in either normal oxygen conditions throughout development (‘N’) or shifted to 5% O_2_ at 120 h AEL (‘H’). n (# of vials of 30 larvae) ≥ 3 per condition.). **(D)** Average time to pupation of larvae, either *spok >* + or *spok > sSpi* (expressing cleaved/secreted spitz), reared in either normal oxygen conditions throughout development (‘N’) or shifted to 5% O_2_ at 120 h AEL (‘H’). n (# of vials of 30 larvae) ≥ 3 per condition. * denotes p < 0.05.

## Discussion

In this paper, we describe a mechanism by which *Drosophila* larvae delay their maturation in hypoxia. We show that hypoxia can slow early larval growth and development to CW but can also act post-CW to suppress ecdysone biosynthesis in the PG. This post-CW effect involves dampening the production of the EGF ligand, spitz, which normally is needed to activate EGF signalling, a major stimulator of ecdysone biosynthesis in the PG.

The delay in attaining CW in hypoxic larvae is similar to that seen when larvae are raised on low-nutrient food. The insulin/TOR signalling pathways are the main pathways that couple dietary nutrients to control tissue and body growth. When nutrients are limited, insulin/TOR signaling is reduced and larval growth is slowed (Boulan *et al*., 2015; Texada *et al*., 2020). The delayed attainment of CW caused by hypoxia exposure in early larval development is most likely due to a similar lowering of systemic insulin/TOR. For example, hypoxia has been shown to promote HIF-1-alpha-independent inhibition of TOR signaling (Lee *et al*., 2019; Texada *et al*., 2019), suppress systemic insulin signaling by reducing insulin secretion from the brain (Texada *et al*., 2019; Wong *et al*., 2014), and upregulate expression of the insulin antagonist, ImpL2 (Kapali *et al*., 2022).

However, an important finding in our work is that hypoxia can also act at the post-CW stage late in larval development to delay maturation by suppressing ecdysone synthesis. This is consistent with a previous report showing that modest hypoxia (10% oxygen) reduced hemolymph ecdysone levels at the end of the larval period (Callier *et al*., 2013). We found that the suppression of ecdysone synthesis genes was equally strong when larvae were either raised in hypoxia from hatching or transferred to hypoxia post-CW, suggesting a specific effect of hypoxia on the late larval events that lead to ecdysone biosynthesis. In this post-CW context, there is a difference between the effects of hypoxia and nutrient restriction on maturation. When starved post-CW, larvae can successfully trigger the late ecdysone pulse and mature to the pupal stage. This is thought to be because they have reached a sufficient mass and level of stored nutrients to support metamorphosis during the non-feeding pupal stage (Rewitz *et al*., 2013; Yamanaka *et al*., 2013). Indeed, critical weight is operationally defined as the point at which starvation does not lead to growth arrest and failure to maturation. The nutrient regulation of this CW checkpoint has been shown to rely on stimulation of insulin/TOR signaling in the PG (Koyama *et al*., 2014; Layalle *et al*., 2008; Ohhara *et al*., 2017), and genetic stimulation of these pathways in the PG is sufficient to reverse the delay in development caused by nutrient limitation. However, we found that activation of TOR in the PG did not accelerate development in hypoxia and hypoxia exposure after CW produced a substantial delay in development. Our findings suggest that larvae use different maturation strategies to cope with nutrient deprivation and low oxygen, which could reflect distinct metabolic strategies in response to fluctuations in these two environmental cues. For example, while starvation leads to lipid depletion (Heier and Kuhnlein, 2018), we previously showed that larvae increase their lipid levels in hypoxia (Lee *et al*., 2019). The hypoxia-induced accumulation in fat body lipids is needed to support subsequent pupal development (Lee *et al*., 2019). Therefore, it is possible that when exposed to hypoxia, larvae need to extend their growth and feeding period to ensure that they accumulate these lipid stores.

Our data point to decreased Spitz expression and suppression of EGFR/Ras/ERK in the PG as the main way that hypoxia delays larval maturation. PTTH/Torso signaling is well established as a trigger for stimulating ERK-dependent ecdysone synthesis in the PG (McBrayer *et al*., 2007; Pan and O’Connor, 2021; Rewitz *et al*., 2009; Shimell *et al*., 2018). However, we saw that larvae with reduced PTTH/Torso signaling hypoxia extend their developmental period, and this extension is greater than that seen with PTTH/Torso inhibition in normoxia, suggesting that hypoxia and PTTH/Torso suppression function independently to delay development. We also found little effect of knockdown of either Alk or Pvr on developmental timing, suggesting that they are unlikely to be involved in the hypoxia-induced delay in development. These results contrast with recent studies that have implicated both RTKs in the control of larval maturation (Juarez-Carreño and Geissmann, 2023; Pan and O’Connor, 2021). These differences may reflect subtle differences in nutrient conditions between our work and these previous studies, especially since the effects of Pvr signalling on developmental timing are regulated by nutrients (Juarez-Carreño and Geissmann, 2023).

How might hypoxia suppress *spitz* mRNA expression? Our data suggest that the effects are independent of HIF-1 alpha/sima, the best-described mediator of hypoxia-induced changes in gene expression. Although previous studies have shown that HIF-1 alpha plays important roles in other *Drosophila* larval tissues to control adaptations to hypoxia (Centanin et al., 2008; Centanin *et al*., 2005; Lavista-Llanos *et al*., 2002; Li *et al*., 2013; Romero *et al*., 2007; Texada *et al*., 2019), our results add to the body of work indicating that HIF-1 alpha-independent mechanisms are also important in controlling organismal adaptations to hypoxia (Barretto *et al*., 2020; Lee *et al*., 2019; Li *et al*., 2013). The upregulation of *spitz* mRNA expression in the PG at the end of the larval period is triggered by an initial pulse of ecdysone signaling through the ecdysone receptor (EcR) (Cruz *et al*., 2020). These nuclear hormone receptors bind DNA and directly control gene expression. Hence, it is possible that hypoxia can antagonize EcR-dependent expression of Spitz. Interestingly, in *C elegans,* hypoxia has been shown to modulate nuclear hormone receptor activity to antagonize EGF/ERK signaling (Maxeiner et al., 2019). Hypoxia could also control PG gene expression via alpha-ketoglutarate-dependent dioxygenases, which include several chromatin modifiers (Baksh and Finley, 2021). These use oxygen in their reaction cycle and are directly regulated by hypoxia (Chakraborty et al., 2019; Hancock et al., 2015). One member of this family of chromatin modifiers, KDM5, functions in the PG to control developmental timing in *Drosophila* (Drelon et al., 2019), while mammalian members of this family have been shown to control the expression of different cytokines and growth factors (Tausendschon et al., 2011; Tian et al., 2019).

Our study has implications for understanding how low oxygen levels can affect development in normal and disease conditions. Hypoxia suppression of EGF signaling is conserved in other organisms (Deygas et al., 2018; Maxeiner *et al*., 2019), and altered expression of EGFR signaling components has been reported both in human populations living in high altitudes (Jha et al., 2016) and in *Drosophila* populations which have undergone multiple generations of selection for hypoxia tolerance (Zhou et al., 2008). Also, hypoxia can modulate steroidogenesis and steroid functions in many animals (Bera et al., 2020; Oyedokun et al., 2023; Wang et al., 2008; Yao et al., 2021), and diseases that can result in reduced systemic oxygen levels in humans - such as chronic obstructive pulmonary disease, sleep apnea or asthma - can impact steroid-induced development and sexual maturation (Drosdzol et al., 2007; Gaudino et al., 2021; Shaw et al., 2013). Thus, the mechanisms that we describe here in *Drosophila* may reflect more general mechanisms by which low oxygen can affect animal development.

## Materials and Methods

### *Drosophila* Food and Genetics

The experimental larvae were raised on food consisting of 150 g agar, 1600 g cornmeal, 770 g Torula yeast, 675 g sucrose, 2340 g D-glucose, and 240 mL of an acid mixture (propionic acid/phosphoric acid) for every 34 L of water, and maintained at 25 °C. For starvation experiments, larvae were switched from standard food to vials containing only agar/water. For all GAL4/UAS experiments, experimental larvae were those obtained from crossing GAL4 lines with the appropriate UAS line. Control animals were obtained by mating the relevant GAL4 line with flies of the same genetic background as the relevant experimental UAS transgene line.

### Drosophila Stocks

The following fly stocks were used in this study: *spok-GAL4;UAS-dicer* (BDSC #80578), *phm-GAL4* (BDSC #80577), *esg-GAL4* (BDSC #84303), *elav-GAL4* (BDSC #458), *UAS-Egfr-RNAi* (BDSC #25781), *P[lacW]spi^s3547^/CyO* (BDSC #10462), *UAS-sSpi* (BDSC #63134), *UAS-spi-RNAi* (VDRC #3920), *UAS-vn-RNAi* (VDRC #50358), *UAS-Rheb* (BDSC#9688), *ywhsflp^122^;+;+* (Prober and Edgar, 2002),*yv* (BDSC #36303), *w^1118^* (BDSC #5905), GD control line (VDRC #60000), *UAS-Raf^GOF^* (Prober and Edgar, 2002), *UAS-sima RNAi* (VDRC #106187), *UAS-sima-RNAi* (*TRiP*) (BDSC #33895), KK control line (VDRC #60100), *w[*]; TI[RFP[3xP3.cUa]=TI]Ptth[attP]* (BDSC #84568), *r4-GAL4* (BDSC #33832), *rn-GAL4* (BDSC #78345) The following stocks were gifts from Mike O’Connor: *P0206*-*GAL4, phm-GAL4;UAS-dicer*, *ptth^120F2A^*, *UAS-λTOP*.

### Hypoxia Exposure

Vials containing larvae were placed in an airtight chamber into which a gas mixture of 5% oxygen and 95% nitrogen was continuously flowed. The rate of flow was controlled using an Aalborg model P gas flow meter. The developmental age of exposure is indicated in individual figures – either 24, 72, 96 or 120 h AEL. For developmental timing experiments, larvae were left in hypoxia to pupate until no further pupation was observed for 24 hours after the initial onset of pupation was observed in all vials. During this time, developmental timing was measured by counting pupae through the transparent walls of the hypoxia chamber.

### Measurement of *Drosophila* Developmental Timing

For each experimental cross, females were allowed to lay eggs on a grape juice/agar plate smeared with yeast paste for four hours. 24hours later, newly hatched larvae were transferred to vials containing our standard lab fly food. 30 larvae were transferred per vial and the number of larvae that pupated in each vial were subsequently counted twice a day. A minimum of three vials of 30 larvae were used to calculate the average time to pupation and percent pupation for each experimental group.

### Pupal Volume Measurements

Pupae were positioned ventral side-down on a petri dish. Images of pupae were captured with the ZEISS SteREO Discovery.V8 microscope and ZEN imaging software (blue edition) at 1.25x magnification. From these images, the ZEN software was used to measure the width and length of each pupa (in µm). These values were then entered into a mathematical formula for the volume of an ellipsoid (4/3π x L/2 x W). Each value resulting from this calculation constituted the pupal volume of one animal.

### Quantitative real-time PCR (qRT-PCR)

Total RNA was extracted from larvae in groups of 8 using TRIzol according to manufacturer instructions (Invitrogen; 15596–018). RNA samples were then DNase-treated with Ambion Turbo DNase according to manufacturer’s instructions (Ambion; 2238 G) and reverse transcribed with Superscript II (Invitrogen; 100004925). Resultant cDNA was used as a template for qRT–PCR (ABI 7500 real time PCR system using SyBr Green PCR mix) with the below primer pairs. PCR data were normalized to levels of Rpl32.

spookier forward: TATCTCTTGGGCACACTCGCTG
spookier reverse: GCCGAGCTAAATTTCTCCGCTT

phantom forward: GGATTTCTTTCGGCGCGATGTG
phantom reverse: TGCCTCAGTATCGAAAAGCCGT

spitz forward: CGCCCAAGAATGAAAGAGAG
spitz reverse: AGGTATGCTGCTGGTGGAAC

rpl32 forward: ATGCTAAGCTGTCGCACAAA
rpl32 reverse: GTTCGATCCGTAACCGATGT

### Starvation treatment

*w^1118^* larvae were collected at 24 h AEL and transferred into regular food vials (30 larvae per vial). At 72, 96 and 120 h AEL, larvae were removed from these vials by suspending them in 20% sucrose and transferred to vials containing either regular food or an equal volume of 0.8% agar.

### Determination of Age at Critical Weight Attainment in Hypoxia and Normoxia

*w^1118^* larvae were collected and transferred into regular food vials (30 larvae per vial). These vials were either placed into hypoxia to develop or left at ambient oxygen. At 96, 100, 112, 116, 120, 124 and 136 h AEL, larvae were transferred to 0.8% agar as in the above protocol and left to develop in ambient oxygen. The resulting percent pupation was calculated and plotted as a function of the age at starvation for each vial. A minimum of 3 vials of 30 larvae were used to calculate the mean percent pupation for each condition at each starvation time point. The point on the x-axis (age of starvation in h AEL) on the graph where >50% mean pupation was achieved was defined as the age at critical weight attainment for each treatment group (normoxia or hypoxia).

### 20-hydroxyecdysone feeding

*w^1118^* larvae were collected at 24 h AEL and transferred into vials containing food supplemented either with 20-hydroxyecdysone (Sigma-Aldrich CAS number 5289-74-7) dissolved in 95% ethanol (final 20E concentration of 0.3 mg/mL) or an equal volume of 95% ethanol alone.

### Immunostaining and Microscopy

Larvae were inverted and fixed in 8% formaldehyde (Electron Microscopy Science, Hatfield, U.S.A.) in PBS at room temperature for 30 minutes, then blocked for 2 hours in 1% BSA in PBS/0.1% Triton-X 100 at room temperature. Blocked samples were then incubated at 4°C overnight in primary antibody (rabbit anti-beta-galactosidase, Jackson ImmunoResearch Laboratories) dissolved at a 1:200 dilution in 1% BSA in PBS/0.1% Triton-X 100. Tissues were then incubated in secondary antibody (Alexa Fluor 568 (Molecular Probes) goat anti-rabbit secondary antibody (1:2000). DNA was visualized using Hoechst 33258 (Invitrogen, 1:10 000). Brain-Ring gland complexes were then dissected out and mounted on glass slides using Vectashield mounting media. Confocal micrographs were captured using a ZEISS confocal microscope LSM 880.

### Quantification of Spitz expression in PG cells

Confocal slices of ring glands from *spitz-LacZ* larvae stained for β-galactosidase were used for analysis in Fiji ImageJ2 (Version 2.3.0). β-galactosidase staining intensity was measured by taking four PG cell nuclei (as discerned by Hoechst staining) at random per PG and measuring the RFP fluorescence intensity in a box of fixed area which fit inside the PG cell nucleus. This way, 4 measurements were taken per animal.

### Statistical Analyses

The mean times to pupation and pupal volume measurements were analyzed by two-way ANOVA followed by unpaired Student’s t-test with Welch’s correction for unequal variance, using an alpha value of 0.05. For qRT-PCR, unpaired Student’s t-test with Welch’s correction for unequal variance was employed for pairwise comparisons. Graphs and statistical analyses were produced using GraphPad Prism 9.0.0 for Windows by GraphPad Software (San Diego, California USA).

---

**Figure S1.**
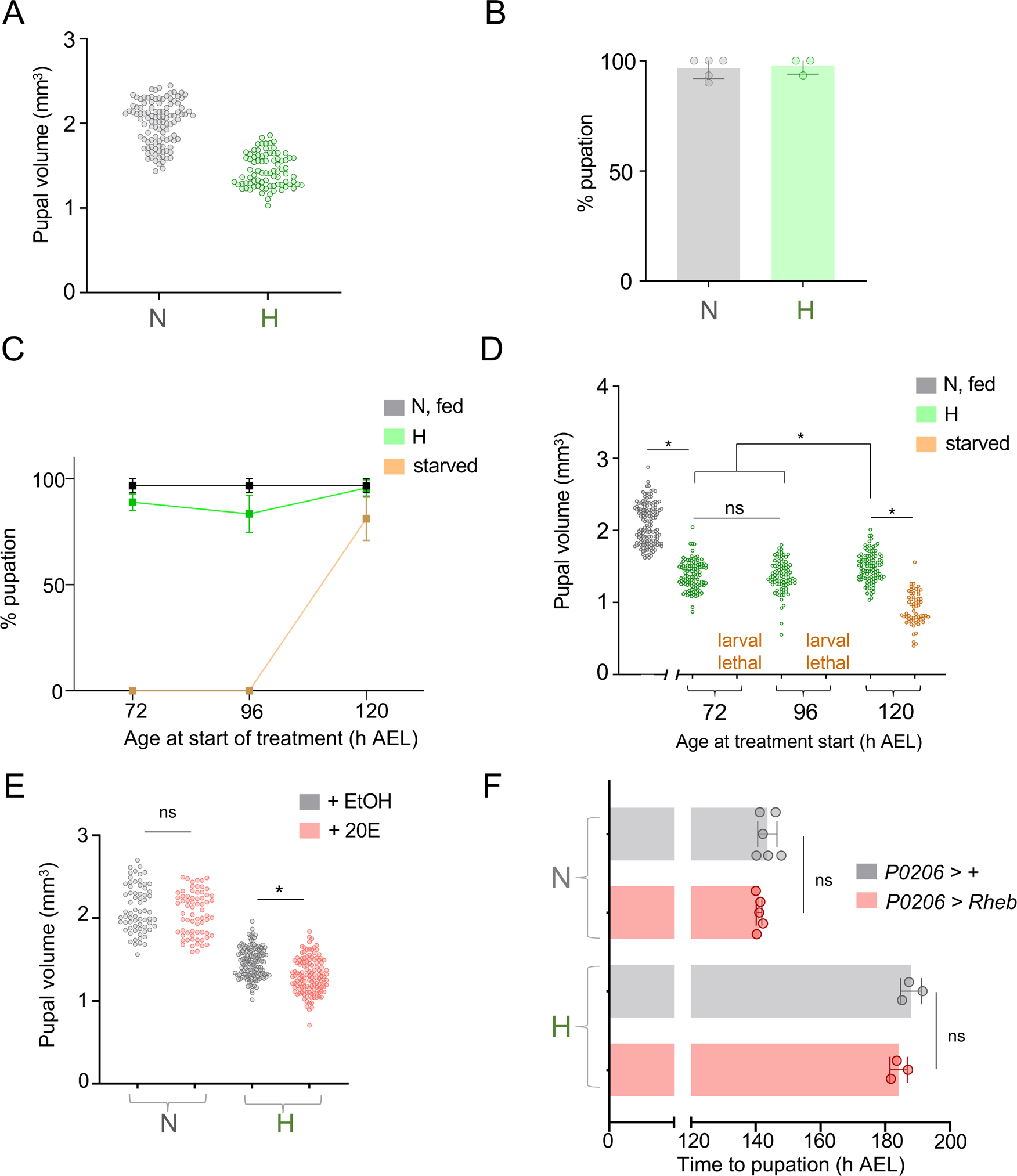
**(related to Figure 1). (A)** Pupal volume of *w^1118^* larvae reared in normoxia throughout development (‘N’) or in hypoxia from 24 h AEL (‘H’). n (# of pupae) = 112 (normoxia) and 83 (hypoxia) **(B)** % survival to the pupal stage of *w^1118^* larvae reared in normoxia throughout development or in hypoxia from 24 h AEL. Each data point represents the average calculated from a vial of 30 larvae. n (# of vials of 30 larvae) ≥ 3 per condition. **(C)** % survival to the pupal stage of *w^1118^ larvae* either starved or exposed to 5% O_2_ at the indicated larval age. Each data point represents the average calculated from a vial of 30 larvae. n (# of vials of 30) ≥ 3 per condition. **(D)** Pupal volume of *w^1118^ larvae* either starved or exposed to 5% O_2_ (‘H’) at the indicated larval age. n (# of pupae) = 148 (normoxia), 107 (H @ 72 h AEL), 91 (H @ 96 h AEL), 107 (H @ 120 h AEL), 62 (starvation @ 120 h AEL). **(E)** Pupal volume of *w^1118^* larvae fed either 20-hydroxyecdysone (20E) or ethanol (EtOH) vehicle, reared in either ambient oxygen or 5% oxygen from 120 h AEL (*i.e.* post-CW). n (# of pupae) = 68 (N, EtOH), 69 (N, 20E), 137 (H, EtOH), 127 (H, 20E). **(F)** Average time to pupation of larvae, either *P0206 >* + or *P0206 > Rheb*, reared in either normal oxygen conditions throughout development or shifted to 5% O_2_ at 120 h AEL. n (# of vials of 30 larvae) ≥ 3 per condition. * denotes p < 0.05.

**Figure S2.**
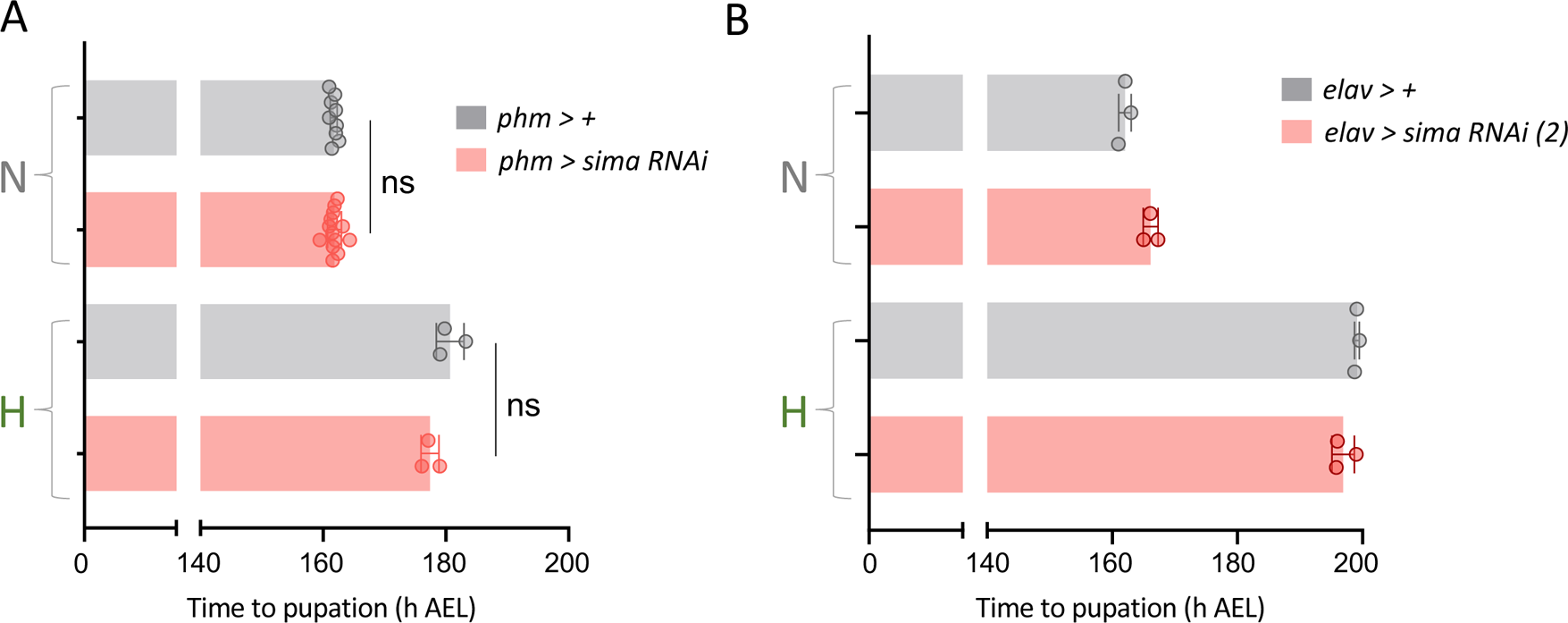
**(related to Figure 2). (A)** Average time to pupation of larvae, either *phm >* + or *phm > sima-RNAi*, reared in either normal oxygen conditions throughout development (‘N’) or shifted to 5% O_2_ at 120 h AEL (‘H’) n (# of vials of 30 larvae) ≥ 3 per condition. **(B)** Average time to pupation of larvae, either *elav >* + or *elav > sima-RNAi (2)*, reared in either normal oxygen conditions throughout development or shifted to 5% O_2_ at 120 h AEL. n (# of vials of 30 larvae) ≥ 3 per condition. * denotes p < 0.05.

**Figure S3.**
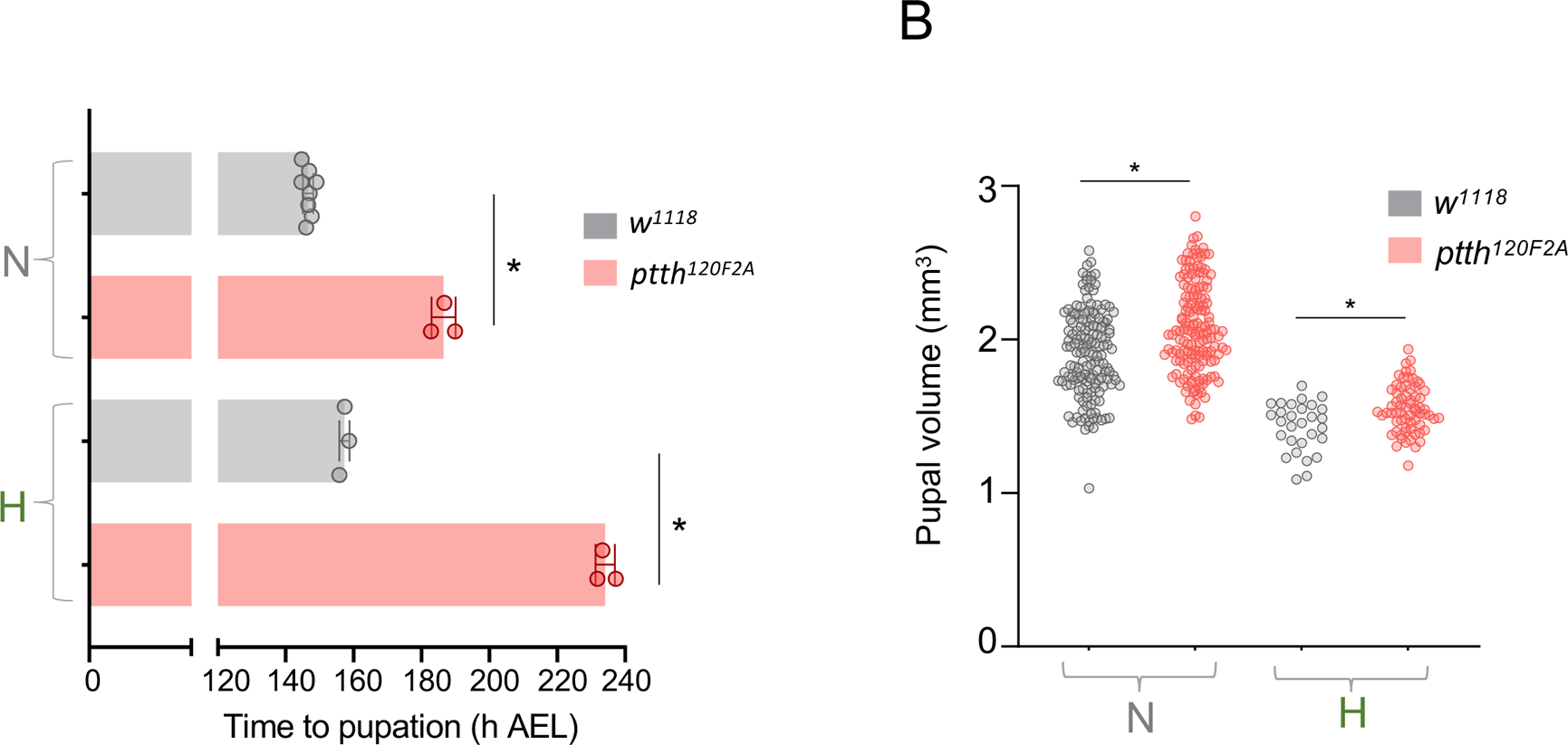
**(related to Figure 3). (A)** Average time to pupation of larvae, either *w^1118^* or *ptth^120F2A^* (an alternative ptth null mutant) reared in either normal oxygen conditions throughout development (‘N’) or shifted to 5% O_2_ at 120 h AEL (‘H’). n (# of vials of 30 larvae) ≥ 3 per condition. **(B)** Pupal size of animals, either *w^1118^* or mutant for *ptth*, reared in normoxia or hypoxia from 120 h AEL. Each data point represents body size measured for one animal. n (# of pupae) = 150 (N *w^1118^*), 150 (N *ptth^120F2A^*), 29 (H *w^1118^*), 67 (H *ptth^120F2A^*). * denotes p < 0.05.

## References

Baksh, S.C., and Finley, L.W.S. (2021). Metabolic Coordination of Cell Fate by alpha-Ketoglutarate-Dependent Dioxygenases. Trends in cell biology 31, 24–36. 10.1016/j.tcb.2020.09.010

Barretto, E.C., Polan, D.M., Beevor-Potts, A.N., Lee, B., and Grewal, S.S. (2020). Tolerance to Hypoxia Is Promoted by FOXO Regulation of the Innate Immunity Transcription Factor NF-kappaB/Relish in Drosophila. Genetics. 10.1534/genetics.120.303219

Bera, A., Chadha, N.K., Dasgupta, S., Chakravarty, S., and Sawant, P.B. (2020). Hypoxia-mediated inhibition of cholesterol synthesis leads to disruption of nocturnal sex steroidogenesis in the gonad of koi carp, Cyprinus carpio. Fish Physiol Biochem 46, 2421–2435. 10.1007/s10695-020-00887-5

Bickler, P.E., and Buck, L.T. (2007). Hypoxia tolerance in reptiles, amphibians, and fishes: life with variable oxygen availability. Annu Rev Physiol 69, 145–170. 10.1146/annurev.physiol.69.031905.162529

Boulan, L., Milan, M., and Leopold, P. (2015). The Systemic Control of Growth. Cold Spring Harb Perspect Biol 7. 10.1101/cshperspect.a019117

Britton, J.S., and Edgar, B.A. (1998). Environmental control of the cell cycle in Drosophila: nutrition activates mitotic and endoreplicative cells by distinct mechanisms. Development 125, 2149–2158.

Callier, V., Hand, S.C., Campbell, J.B., Biddulph, T., and Harrison, J.F. (2015). Developmental changes in hypoxic exposure and responses to anoxia in Drosophila melanogaster. J Exp Biol 218, 2927–2934. 10.1242/jeb.125849

Callier, V., and Nijhout, H.F. (2014). Plasticity of insect body size in response to oxygen: integrating molecular and physiological mechanisms. Curr Opin Insect Sci 1, 59–65. 10.1016/j.cois.2014.05.007

Callier, V., Shingleton, A.W., Brent, C.S., Ghosh, S.M., Kim, J., and Harrison, J.F. (2013). The role of reduced oxygen in the developmental physiology of growth and metamorphosis initiation in Drosophila melanogaster. J Exp Biol 216, 4334–4340. 10.1242/jeb.093120

Centanin, L., Dekanty, A., Romero, N., Irisarri, M., Gorr, T.A., and Wappner, P. (2008). Cell autonomy of HIF effects in Drosophila: tracheal cells sense hypoxia and induce terminal branch sprouting. Developmental cell 14, 547–558. 10.1016/j.devcel.2008.01.020

Centanin, L., Ratcliffe, P.J., and Wappner, P. (2005). Reversion of lethality and growth defects in Fatiga oxygen-sensor mutant flies by loss of hypoxia-inducible factor-alpha/Sima. EMBO Rep 6, 1070–1075. 10.1038/sj.embor.7400528

Chakraborty, A.A., Laukka, T., Myllykoski, M., Ringel, A.E., Booker, M.A., Tolstorukov, M.Y., Meng, Y.J., Meier, S.R., Jennings, R.B., Creech, A.L., et al. (2019). Histone demethylase KDM6A directly senses oxygen to control chromatin and cell fate. Science (New York, N.Y.) 363, 1217–1222. 10.1126/science.aaw1026

Church, R.B., and Robertson, F.W. (1966). Biochemical analysis of genetic differences in the growth of Drosophila. Genet Res 7, 383–407.

Colombani, J., Bianchini, L., Layalle, S., Pondeville, E., Dauphin-Villemant, C., Antoniewski, C., Carre, C., Noselli, S., and Leopold, P. (2005). Antagonistic actions of ecdysone and insulins determine final size in Drosophila. Science (New York, N.Y.) 310, 667–670. 10.1126/science.1119432

Cruz, J., Martin, D., and Franch-Marro, X. (2020). Egfr Signaling Is a Major Regulator of Ecdysone Biosynthesis in the Drosophila Prothoracic Gland. Current biology : CB 30, 1547–1554 e1544. 10.1016/j.cub.2020.01.092

Danielsen, E.T., Moeller, M.E., and Rewitz, K.F. (2013). Nutrient signaling and developmental timing of maturation. Curr Top Dev Biol 105, 37–67. 10.1016/B978-0-12-396968-2.00002-6

Deygas, M., Gadet, R., Gillet, G., Rimokh, R., Gonzalo, P., and Mikaelian, I. (2018). Redox regulation of EGFR steers migration of hypoxic mammary cells towards oxygen. Nat Commun 9, 4545. 10.1038/s41467-018-06988-3

Di Cara, F., and King-Jones, K. (2016). The Circadian Clock Is a Key Driver of Steroid Hormone Production in Drosophila. Current biology : CB 26, 2469–2477. 10.1016/j.cub.2016.07.004

Drelon, C., Rogers, M.F., Belalcazar, H.M., and Secombe, J. (2019). The histone demethylase KDM5 controls developmental timing in Drosophila by promoting prothoracic gland endocycles. Development 146. 10.1242/dev.182568

Drosdzol, A., Skrzypulec, V., Wilk, K., and Nowosielski, K. (2007). The influence of bronchial asthma on sexual maturation of girls. J Physiol Pharmacol 58 Suppl 5, 155–163.

Ducsay, C.A., Goyal, R., Pearce, W.J., Wilson, S., Hu, X.Q., and Zhang, L. (2018). Gestational Hypoxia and Developmental Plasticity. Physiol Rev 98, 1241–1334. 10.1152/physrev.00043.2017

Farzin, M., Albert, T., Pierce, N., VandenBrooks, J.M., Dodge, T., and Harrison, J.F. (2014). Acute and chronic effects of atmospheric oxygen on the feeding behavior of Drosophila melanogaster larvae. J Insect Physiol 68, 23–29. 10.1016/j.jinsphys.2014.06.017

Gaudino, R., Dal Ben, S., Cavarzere, P., Volpi, S., Piona, C., Boner, A., Antoniazzi, F., and Piacentini, G. (2021). Delayed age at menarche in chronic respiratory diseases. Eur J Clin Invest 51, e13461. 10.1111/eci.13461

Ghosh, S., Leng, W., Wilsch-Brauninger, M., Barrera-Velazquez, M., Leopold, P., and Eaton, S. (2022). A local insulin reservoir in Drosophila alpha cell homologs ensures developmental progression under nutrient shortage. Current biology : CB 32, 1788–1797 e1785. 10.1016/j.cub.2022.02.068

Grewal, S.S. (2009). Insulin/TOR signaling in growth and homeostasis: a view from the fly world. Int J Biochem Cell Biol 41, 1006–1010. 10.1016/j.biocel.2008.10.010

Hancock, R.L., Dunne, K., Walport, L.J., Flashman, E., and Kawamura, A. (2015). Epigenetic regulation by histone demethylases in hypoxia. Epigenomics 7, 791–811. 10.2217/epi.15.24

Harrison, J.F., Greenlee, K.J., and Verberk, W. (2018). Functional Hypoxia in Insects: Definition, Assessment, and Consequences for Physiology, Ecology, and Evolution. Annu Rev Entomol 63, 303–325. 10.1146/annurev-ento-020117-043145

Harrison, J.F., and Haddad, G.G. (2011). Effects of oxygen on growth and size: synthesis of molecular, organismal, and evolutionary studies with Drosophila melanogaster. Annu Rev Physiol 73, 95–113. 10.1146/annurev-physiol-012110-142155

Heier, C., and Kuhnlein, R.P. (2018). Triacylglycerol Metabolism in Drosophila melanogaster. Genetics 210, 1163–1184. 10.1534/genetics.118.301583

Hietakangas, V., and Cohen, S.M. (2009). Regulation of tissue growth through nutrient sensing. Annual review of genetics 43, 389–410. 10.1146/annurev-genet-102108-134815

Jha, A.R., Zhou, D., Brown, C.D., Kreitman, M., Haddad, G.G., and White, K.P. (2016). Shared Genetic Signals of Hypoxia Adaptation in Drosophila and in High-Altitude Human Populations. Mol Biol Evol 33, 501–517. 10.1093/molbev/msv248

Juarez-Carreño, S., and Geissmann, F. (2023). The macrophage genetic cassette inr/dtor/pvf2 is a nutritional status checkpoint for developmental timing. bioRxiv, 2023.2001.2005.522883. 10.1101/2023.01.05.522883

Kannangara, J.R., Mirth, C.K., and Warr, C.G. (2021). Regulation of ecdysone production in Drosophila by neuropeptides and peptide hormones. Open Biol 11, 200373. 10.1098/rsob.200373

Kapali, G.P., Callier, V., Gascoigne, S.J.L., Harrison, J.F., and Shingleton, A.W. (2022). The steroid hormone ecdysone regulates growth rate in response to oxygen availability. Sci Rep 12, 4730. 10.1038/s41598-022-08563-9

Koyama, T., and Mirth, C.K. (2018). Unravelling the diversity of mechanisms through which nutrition regulates body size in insects. Curr Opin Insect Sci 25, 1–8. 10.1016/j.cois.2017.11.002

Koyama, T., Rodrigues, M.A., Athanasiadis, A., Shingleton, A.W., and Mirth, C.K. (2014). Nutritional control of body size through FoxO-Ultraspiracle mediated ecdysone biosynthesis. eLife 3. 10.7554/eLife.03091

Koyama, T., Texada, M.J., Halberg, K.A., and Rewitz, K. (2020). Metabolism and growth adaptation to environmental conditions in Drosophila. Cell Mol Life Sci 77, 4523–4551. 10.1007/s00018-020-03547-2

Lavista-Llanos, S., Centanin, L., Irisarri, M., Russo, D.M., Gleadle, J.M., Bocca, S.N., Muzzopappa, M., Ratcliffe, P.J., and Wappner, P. (2002). Control of the hypoxic response in Drosophila melanogaster by the basic helix-loop-helix PAS protein similar. Mol Cell Biol 22, 6842–6853.

Layalle, S., Arquier, N., and Leopold, P. (2008). The TOR pathway couples nutrition and developmental timing in Drosophila. Developmental cell 15, 568–577. 10.1016/j.devcel.2008.08.003

Lee, B., Barretto, E.C., and Grewal, S.S. (2019). TORC1 modulation in adipose tissue is required for organismal adaptation to hypoxia in Drosophila. Nat Commun 10, 1878. 10.1038/s41467-019-09643-7

Li, Y., Padmanabha, D., Gentile, L.B., Dumur, C.I., Beckstead, R.B., and Baker, K.D. (2013). HIF- and non-HIF-regulated hypoxic responses require the estrogen-related receptor in Drosophila melanogaster. PLoS genetics 9, e1003230. 10.1371/journal.pgen.1003230

Malita, A., and Rewitz, K. (2021). Interorgan communication in the control of metamorphosis. Curr Opin Insect Sci 43, 54–62. 10.1016/j.cois.2020.10.005

Markow, T.A. (2015). The secret lives of Drosophila flies. eLife 4. 10.7554/eLife.06793

Maxeiner, S., Grolleman, J., Schmid, T., Kammenga, J., and Hajnal, A. (2019). The hypoxia-response pathway modulates RAS/MAPK-mediated cell fate decisions in Caenorhabditis elegans. Life Sci Alliance 2. 10.26508/lsa.201800255

McBrayer, Z., Ono, H., Shimell, M., Parvy, J.P., Beckstead, R.B., Warren, J.T., Thummel, C.S., Dauphin-Villemant, C., Gilbert, L.I., and O’Connor, M.B. (2007). Prothoracicotropic hormone regulates developmental timing and body size in Drosophila. Developmental cell 13, 857–871. 10.1016/j.devcel.2007.11.003

Mirth, C., Truman, J.W., and Riddiford, L.M. (2005). The role of the prothoracic gland in determining critical weight for metamorphosis in Drosophila melanogaster. Current biology : CB 15, 1796–1807. 10.1016/j.cub.2005.09.017

Mirth, C.K., Saunders, T.E., and Amourda, C. (2021). Growing Up in a Changing World: Environmental Regulation of Development in Insects. Annu Rev Entomol 66, 81–99. 10.1146/annurev-ento-041620-083838

Moore, L.G., Charles, S.M., and Julian, C.G. (2011). Humans at high altitude: hypoxia and fetal growth. Respir Physiol Neurobiol 178, 181–190. 10.1016/j.resp.2011.04.017

Ohhara, Y., Kobayashi, S., and Yamanaka, N. (2017). Nutrient-Dependent Endocycling in Steroidogenic Tissue Dictates Timing of Metamorphosis in Drosophila melanogaster. PLoS genetics 13, e1006583. 10.1371/journal.pgen.1006583

Oyedokun, P.A., Akhigbe, R.E., Ajayi, L.O., and Ajayi, A.F. (2023). Impact of hypoxia on male reproductive functions. Mol Cell Biochem 478, 875–885. 10.1007/s11010-022-04559-1

Pan, X., Connacher, R.P., and O’Connor, M.B. (2021). Control of the insect metamorphic transition by ecdysteroid production and secretion. Curr Opin Insect Sci 43, 11–20. 10.1016/j.cois.2020.09.004

Pan, X., and O’Connor, M.B. (2021). Coordination among multiple receptor tyrosine kinase signals controls Drosophila developmental timing and body size. Cell reports 36, 109644. 10.1016/j.celrep.2021.109644

Peck, L.S., and Maddrell, S.H. (2005). Limitation of size by hypoxia in the fruit fly Drosophila melanogaster. J Exp Zool A Comp Exp Biol 303, 968–975. 10.1002/jez.a.211

Prober, D.A., and Edgar, B.A. (2002). Interactions between Ras1, dMyc, and dPI3K signaling in the developing Drosophila wing. Genes Dev 16, 2286–2299.

Rewitz, K.F., Yamanaka, N., Gilbert, L.I., and O’Connor, M.B. (2009). The insect neuropeptide PTTH activates receptor tyrosine kinase torso to initiate metamorphosis. Science (New York, N.Y.) 326, 1403–1405. 10.1126/science.1176450

Rewitz, K.F., Yamanaka, N., and O’Connor, M.B. (2013). Developmental checkpoints and feedback circuits time insect maturation. Curr Top Dev Biol 103, 1–33. 10.1016/B978-0-12-385979-2.00001-0

Romero, N.M., Dekanty, A., and Wappner, P. (2007). Cellular and developmental adaptations to hypoxia: a Drosophila perspective. Methods Enzymol 435, 123–144. 10.1016/S0076-6879(07)35007-6

Semenza, G.L. (2011). Oxygen sensing, homeostasis, and disease. N Engl J Med 365, 537–547. 10.1056/NEJMra1011165

Shaw, N.D., Goodwin, J.L., Silva, G.E., Hall, J.E., Quan, S.F., and Malhotra, A. (2013). Obstructive sleep apnea (OSA) in preadolescent girls is associated with delayed breast development compared to girls without OSA. J Clin Sleep Med 9, 813–818. 10.5664/jcsm.2928

Shimada-Niwa, Y., and Niwa, R. (2014). Serotonergic neurons respond to nutrients and regulate the timing of steroid hormone biosynthesis in Drosophila. Nat Commun 5, 5778. 10.1038/ncomms6778

Shimell, M., Pan, X., Martin, F.A., Ghosh, A.C., Leopold, P., O’Connor, M.B., and Romero, N.M. (2018). Prothoracicotropic hormone modulates environmental adaptive plasticity through the control of developmental timing. Development 145. 10.1242/dev.159699

Tausendschon, M., Dehne, N., and Brune, B. (2011). Hypoxia causes epigenetic gene regulation in macrophages by attenuating Jumonji histone demethylase activity. Cytokine 53, 256–262. 10.1016/j.cyto.2010.11.002

Texada, M.J., Jorgensen, A.F., Christensen, C.F., Koyama, T., Malita, A., Smith, D.K., Marple, D.F.M., Danielsen, E.T., Petersen, S.K., Hansen, J.L., et al. (2019). A fat-tissue sensor couples growth to oxygen availability by remotely controlling insulin secretion. Nat Commun 10, 1955. 10.1038/s41467-019-09943-y

Texada, M.J., Koyama, T., and Rewitz, K. (2020). Regulation of Body Size and Growth Control. Genetics 216, 269–313. 10.1534/genetics.120.303095

Tian, Z., Yao, L., Shen, Y., Guo, X., and Duan, X. (2019). Histone H3K9 demethylase JMJD1A is a co-activator of erythropoietin expression under hypoxia. Int J Biochem Cell Biol 109, 33–39. 10.1016/j.biocel.2019.01.022

Troha, K., and Ayres, J.S. (2020). Metabolic Adaptations to Infections at the Organismal Level. Trends Immunol 41, 113–125. 10.1016/j.it.2019.12.001

Wang, S., Yuen, S.S., Randall, D.J., Hung, C.Y., Tsui, T.K., Poon, W.L., Lai, J.C., Zhang, Y., and Lin, H. (2008). Hypoxia inhibits fish spawning via LH-dependent final oocyte maturation. Comp Biochem Physiol C Toxicol Pharmacol 148, 363–369. 10.1016/j.cbpc.2008.03.014

Wang, T., Hung, C.C., and Randall, D.J. (2006). The comparative physiology of food deprivation: from feast to famine. Annu Rev Physiol 68, 223–251. 10.1146/annurev.physiol.68.040104.105739

Wong, D.M., Shen, Z., Owyang, K.E., and Martinez-Agosto, J.A. (2014). Insulin- and warts-dependent regulation of tracheal plasticity modulates systemic larval growth during hypoxia in Drosophila melanogaster. PLoS One 9, e115297. 10.1371/journal.pone.0115297

Yamanaka, N., Rewitz, K.F., and O’Connor, M.B. (2013). Ecdysone control of developmental transitions: lessons from Drosophila research. Annu Rev Entomol 58, 497–516. 10.1146/annurev-ento-120811-153608

Yao, S., Lopez-Tello, J., and Sferruzzi-Perri, A.N. (2021). Developmental programming of the female reproductive system-a review. Biol Reprod 104, 745–770. 10.1093/biolre/ioaa232

Zeng, J., Huynh, N., Phelps, B., and King-Jones, K. (2020). Snail synchronizes endocycling in a TOR-dependent manner to coordinate entry and escape from endoreplication pausing during the Drosophila critical weight checkpoint. PLoS Biol 18, e3000609. 10.1371/journal.pbio.3000609

Zhou, D., and Haddad, G.G. (2013). Genetic analysis of hypoxia tolerance and susceptibility in Drosophila and humans. Annu Rev Genomics Hum Genet 14, 25–43. 10.1146/annurev-genom-091212-153439

Zhou, D., Xue, J., Lai, J.C., Schork, N.J., White, K.P., and Haddad, G.G. (2008). Mechanisms underlying hypoxia tolerance in Drosophila melanogaster: hairy as a metabolic switch. PLoS genetics 4, e1000221. 10.1371/journal.pgen.1000221

